# Disrupted Cerebral Peri-Microvascular Glycogen Promotes Capillary Constrictions and Aggravates Ischemia in Mice

**DOI:** 10.1101/2022.08.24.505172

**Authors:** Gokhan Uruk, Sinem Yilmaz-Ozcan, Canan Cakir-Aktas, Aslihan Taskiran-Sag, Buket Donmez-Demir, Jordi Duran, Joan J. Guinovart, Hulya Karatas-Kurşun, Turgay Dalkara, Muge Yemisci

## Abstract

Ischemic stroke results in sudden blood flow cessation, thus, unmet energy requirements. Although the clotted artery can be recanalized and blood flow is restored, brain perfusion may not be fully attained due to microvascular constrictions. Under glucose deprived and hypoxic conditions, glucose derived from the glycogen stored around peri-microvascular astrocyte end-feet may serve as an emergency fuel to meet the metabolic demand during acute period of ischemic stroke. To elucidate the impact of glycogen utilization on brain microcirculation, we administered glycogen phosphorylase inhibitor 1,4-dideoxy-1,4-imino-d-arabinitol (DAB) intracerebroventricularly. Transgenic mice in which glycogen synthase-1 expression was selectively knocked out in central nervous system (GYS1^Nestin-KO^) were also used. Both approaches caused microvascular constrictions mediated by CD13-positive pericyte contractions. When mice with disrupted glycogen utilization were subjected to MCA ischemia, pericyte-mediated microvascular constrictions and the infarct volumes were further increased compared to untreated controls or wild type littermates. Perimicrovascular glycogen depletions were highly correlated with microvascular constrictions as shown by Periodic acid Schiff (PAS) staining and immunolabeling with anti-glycogen antibodies. Imaging of regional cortical blood flow changes during ischemia disclosed severely compromised blood flow dynamics in mice with disrupted glycogen metabolism. In conclusion, disrupting glycogen utilization causes ischemic-like microvascular constrictions under non-ischemic circumstances and increases susceptibility to brain ischemia. Understanding the role of glycogen at neurogliovascular level in brain may provide novel insight to the pathophysiology of ischemic stroke and therapeutic opportunities.

**Graphical Abstract:** 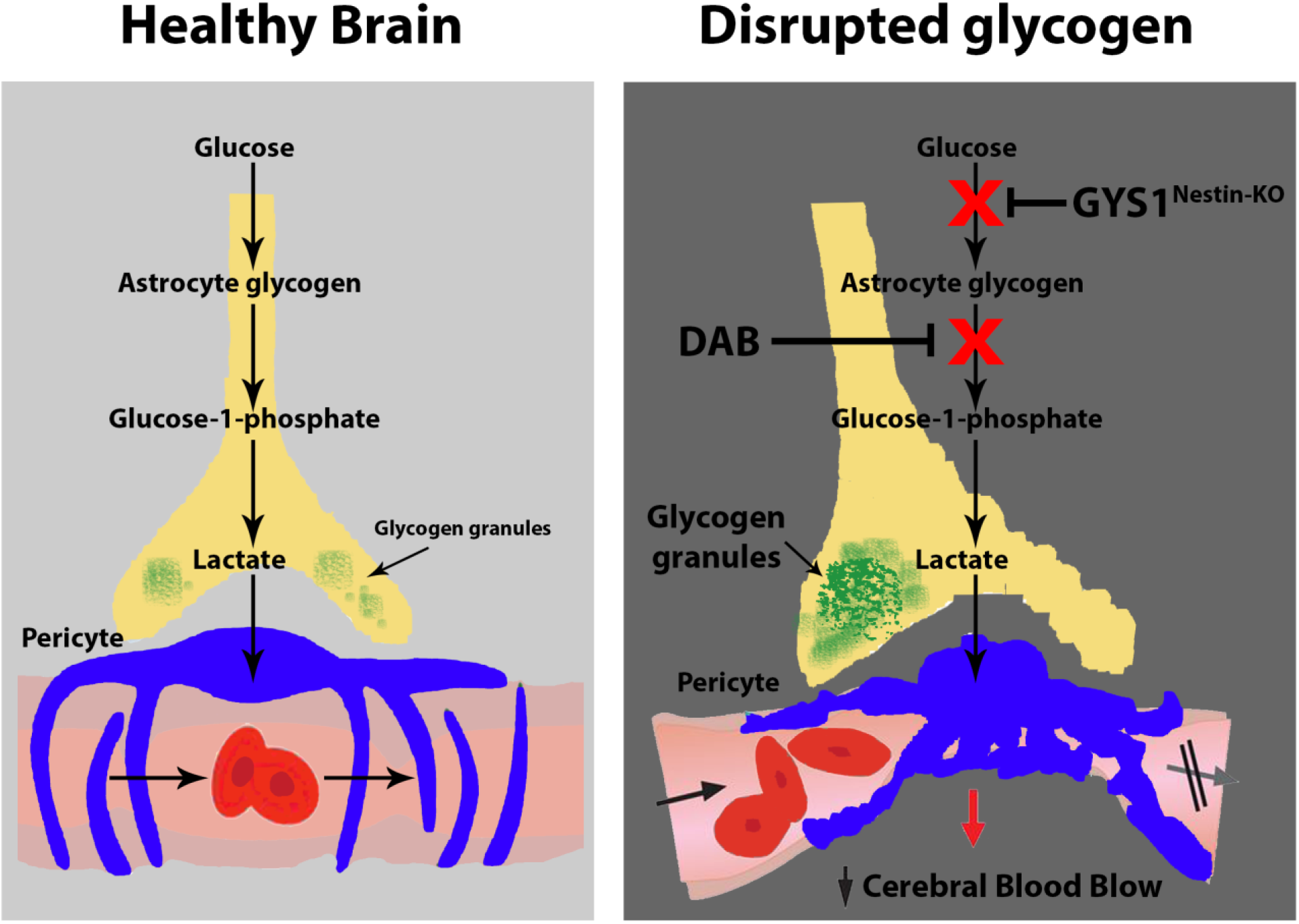

## Introduction

The human brain is comprised of 2% of the body’s weight, however accounts for 20-25% of its resting oxygen metabolism, and glucose consumption (1, 2). Neurons exclusively rely on constant glucose flux and harbor negligible amount of glycogen, as readily usable glucose store. However, glial cells contain glycogen, and although the amount is detected less compared to other tissues, this energy reserve could support the metabolism for a short period of time when needed (3–5). Ischemic stroke results in an abrupt oxygen and glucose insufficiency to the glial cells and neurons thus drastic metabolic changes (5–9). During ischemia, neurons cannot obtain glucose from blood and the glycogen stored in the cells cannot meet the energy requirements more than seconds (10). Therefore, glial cells especially astrocytes play a key role in supporting neurons also under ischemic conditions (11–13). Glial glycogen reserve can metabolically support the tissue by switching glycolysis from blood glucose to glycogen-derived glucose (14). The steady and dynamic balance between glycogenesis and glycogenolysis provides the endogenous energy reserve in ischemia (15).

Patency of the microvasculature is essential for reperfusion, hence survival in cerebral ischemia (16–18). Our previous studies demonstrated that microvascular constrictions due to ischemia are spatially correlated with contracted pericytes in the brain and the retina (17, 19). Although the pathophysiological mechanisms underlying the control of microvascular functions are not fully understood in the brain yet, pericytes are shown to have a role in mediating microvascular constrictions in metabolically compromised retina (1, 17–21).

Glycogen metabolism is tightly regulated in each cell (3, 22, 23). Glycogen synthase is the rate-limiting enzyme in adding glucose moieties to existing glycogen granules thus glycogenesis, and Glycogen phosphorylase (GP) is the key enzyme in glycogen breakdown, glycogenolysis (24). In this study we aimed to investigate the effects of pharmacologically or genetically disrupted peri-microvascular glycogen and superimposed permanent cerebral ischemia on microvasculature. To disrupt glycogen utilization pharmacologically, we intracerebroventricularly administered 1,4-dideoxy-1,4-imino-d-arabinitol (DAB) which potently inhibits glycogen phosphorylase enzyme activity and prevents the use of glycogen (25). For genetic disruption we used central nervous system specific Glycogen Synthase-1 knockout (GYS1^Nestin-KO^) mice (26). We used a permanent cerebral ischemia model to mimic the clinics of the ischemic stroke patients without recanalization (27).

We found that disruption of glycogen metabolism either by i.c.v. DAB administration to mice or in GYS1^Nestin-KO^ mice, induced microvascular constrictions near CD13-positive pericytes as observed in ischemia. Interestingly, when permanent cerebral ischemia was introduced to mice with already disturbed glycogen utilization; CD13-positive pericyte constrictions, and the infarct volume were increased compared to wild type. When L-lactate is administrated i.c.v. as an energy source to bypass inhibited glycogen phosphorylase (25, 28, 29), the DAB-induced CD13-positive pericyte constrictions were diminished. Disruption of glycogen utilization causes energy deficiency in the brain resembling ischemia and leads to increased microvascular constrictions. As we previously observed in mice retina, these microvascular constrictions suggest a role for CD13-positive capillary pericyte contractions, which spatially correspond to ischemia induced peri-microvascular glycogen depletion (19). We suggest that insufficient peri-microvascular glycogen may have a role in the pathophysiology of incomplete reperfusion after cerebral ischemia.

## Materials and Methods

### Animals and Study Approval

The experiments were performed in adult (20–30 g) male and female Swiss albino (n=3/group), C57Bl/6J (WT) (n=3/group) and GYS1^Nestin-KO^ (n=3/group) mice, which were housed under diurnal lighting conditions (12 h light and 12 h darkness) at room temperature (22±2°C, 50-60% humidity). The animals had ad libitum access to water and food. Initial breeding pairs of GYS1^Nestin-KO^ mice ((26)) were obtained from Dr. Duran and Dr. Guinovart at IRB Barcelona and the colony was further expanded, genotyped and maintained at Hacettepe University Institute of Neurological Sciences and Psychiatry. Animal housing, care, and the experimental procedures were all done in accordance with institutional guidelines. The study was approved by Hacettepe University Animal Experimentations Local Ethics Board (Registration number: 2016-60/3 and 2018-56/06).

For all the surgical procedures mentioned below, mice were anesthetized with isoflurane (4-5% induction dose, followed by 1.5-2% maintenance). Corneal reflex and hind-paw nociceptive reflex sensitivity were assessed periodically for depth of anesthesia. O_2_ flow rate was maintained around 2 L/min via a facemask for the duration of experiment to maintain tissue O2 saturation above 92% to avoid anesthesia-induced hypoxia. Body temperature was monitored with a rectal probe and maintained at 37.0 ± 0.2°C by a homeothermic blanket control unit (Harvard Apparatus, USA). Pulse rate and oxygen saturation were monitored with an oximeter (The LifeSense^®^ VET Pulse Oximeter, Nonin Medical Inc., USA) from the right lower limb throughout the experiment. Mice were placed on the heating blanket until fully recovered from anesthesia. The animals which developed hemodynamic instability during experimental procedures were excluded from the study.

### Intracerebroventricular (i.c.v.) Injection

Mice were placed in a rodent stereotaxic frame (WPI, USA) in prone position under isoflurane. The cranium was exposed with a vertical skin incision and a burr hole was opened on either hemisphere. 10 μL 26-gauge Hamilton syringe was placed reaching the ventricle (coordinates respective to Bregma: −0.15 mm posterior, 0.7 mm lateral, 3.15 mm of depth) (30) under surgical stereomicroscope (Nikon SMZ 1000, Nikon). To minimize the tissue trauma due to injection, syringe was introduced in 10 minutes. After waiting for 5 minutes, either DAB alone (n=3, 0.75 μL of 0.25 M in saline), or a combination of DAB (0.25 M) and D-lactate (0.5 M in saline) (n=3) or DAB (0.25 M) and L-lactate (0.5 M in saline) (n=3) were injected i.c.v. in 5 minutes with a pace of 0.15 μL/min (25, 29). Following another 5 minutes wait after the i.c.v. injections, the syringe was removed within 10 minutes. Mid-line incision was sutured via 4-0 nylon suture. Prilocaine was injected subcutaneously to provide pain relief. Mice were placed on the heating blanket until they were fully recovered from anesthesia.

### Thrombotic MCA occlusion (MCAo)

Thrombotic MCAo was performed as previously described (31). Mice were placed in a stereotaxic frame (WPI, USA), the scalp was opened, and the cranial sutures and bregma were exposed. The right temporal muscle was bluntly dissected until the squamous part of the temporal bone was exposed. The area just above the junction between the zygomatic arch and the squamous bone was thinned using a high-speed drill and cooled with saline. The trace of MCA was visualized through the thinned temporal bone. Thin bony film was lifted up by forceps to open up the Burr hole. After obtaining a stable 10-minute epoch of the preischemic regional cerebral blood flow (rCBF) via laser-speckle flowmetry (LSF), 30% FeCl3-saturated filter paper (0.3×1 mm) was placed over the MCA. Filter-paper was removed after 3-10 minutes of placement on observing the cessation of rCBF due to successful clot formation (Karatas et al JCBFM 2011). MCAo was performed for 2 hours and rCBF was continuously monitored during the occlusion and 115 minutes after.

### Laser Speckle Contrast Imaging (LSCI) and Image processing

Laser speckle flowmetry was used to detect the cortical blood flow changes observed through the cranial window as described previously (32). A 785 nm-laser diode (Thorlabs, USA) was used to diffusely illuminate the cortical surface through thinned skull. A CCD camera (Basler 602F, Basler Vision Technologies, Ahrensburg, Germany) attached to a stereomicroscope (Nikon SMZ 1000, Nikon) and custom-developed software (courtesy of A.K. Dunn) was used for LSC which were captured every 10 seconds during 2-hour experiment to construct cortical blood flow maps. A baseline image was constructed by averaging 10 consecutive images before ischemia for each saved inverse correlation time (ICT) image sequence. The field of view was adjusted using a variable magnification objective on the microscope (Nikon SMZ 1000, Nikon). Drift in z-axis was inevitable due to the blood volume changes in the ischemic tissue. To compare the rCBF changes induced by MCAo in selected regions of cortex, LSC images were later imported to ImageJ v1.42q NIH as image sequences and then saved in tiff format for further analysis. To transform contrast values of each image within the saved sequence to CBF values, inverse correlation time images were obtained with MATLAB^®^ (Mathworks) software by taking the camera exposure time (5 ms) into consideration. Subsequent ICT images were differentially divided by the baseline image to yield the relative CBF values and then all calculated values for each pixel were averaged separately for the duration of ischemia. The results were constructed as a single image and pseudo-colored to illustrate the average percent flow change for each pixel during ischemia (ΔCBFisch). To compare rCBF changes between DAB and saline, GYS-1^NestinKO^ and WT mice, two ROIs depicting ischemic core and penumbra within the MCA territory were selected for each experiment from areas devoid of pial vessels. The mean and standard errors of each ROI from ischemic core and peri-infarct area were calculated and histograms were generated with MATLAB software.

### Non-invasive Systolic Blood Pressure Measurement

Besides CBF assessment, systolic blood pressure was measured from the tail via laser-speckle flowmetry. Blood pressure cuff (ADInstruments) was placed to the proximal part of the tail and this cuff was inflated periodically through non-invasive blood pressure (NIBP ML125, ADInstruments) controller. Peanut oil was applied to tail to enhance the reflectance. LSC images were captured every 1 second during the tail cuff inflation for a minute. A baseline image was constructed by averaging 10 consecutive images before inflating the cuff. LSC images were later imported to ImageJ v1.42q NIH as image sequences and then saved in tiff format for further analysis. Two ROIs depicting either of tail veins and artery were selected for each experiment and the graphs regarding blood flow measurements from the tail vessels were constructed. For each measurement, the graphs plotting the tail cuff pressure and tail blood flow values were superimposed. Systolic blood pressure values were obtained by measuring the pressure values at the points where the blood flow was interrupted and restored back. The measurements were performed 2 times 10 minutes before and 120 minutes after MCAo experiments.

### Immunohistochemical Studies

Mice were sacrificed via transcardial perfusion with 4% PFA solution (dissolved in phosphate-buffered saline (PBS)) under lethal dose of chloralhydrate (500 mg/kg intraperitoneally). Harvested brain tissues were post-fixated in PFA solution for 24 hours and then placed in 30% sucrose (in PBS) for at least 2 days to obtain tissue cryopreversation. Brains were cut via a sliding microtome (Lecia SM2000R) with thickness of 50 μm. To permeabilize, the sections were incubated in TBS containing 0.3% Triton-X for 30 minutes at room temperature, then 10 mM sodium citrate (pH: 6.0) for 30 minutes heated to 80 °C for antigen retrieval. The sections were blocked in 1% bovine serum albumin containing 0.3% Triton-X, 10% normal goat serum (NGS) (0.3% TBS-T / 1% BSA, 10% NGS) solution for 1 hour at room temperature and then incubated with primary antibodies (anti-glycogen antibodies ESG1A9 and IV58B6 (courtesy of Dr. Hitoshi Ashida and Dr. Otto Baba), CD13 (Acris Antibodies GmBH, AM26636AF-N) for pericytes) in 0.3% TBS-T overnight at + 4°C. After rinsing, the sections were then incubated with respective secondary antibodies (Alexa Fluor 488 conjugated goat anti-Mouse IgM (Jackson ImmunoResearch, West Grove, PA) for anti-glycogen antibodies, Cy3 conjugated anti-rat IgG (Molecular Probes, ThermoFisher Scientific) for CD13 in 0.3% TBS-T for 90 minutes at room temperature. After rinsing the sections three times for 10 minutes with TBS, sections were then incubated with ‘Fluorescein’ labeled Lycopersicon Esculentum Lectin (dissolved in Tris-buffered saline (TBS); Vector Laboratories, Burlingame, CA) at + 4°C overnight to visualize vessels. Finally, the sections were mounted and covered with anti-fade reagent containing 1/1000 Hoechst 33258 (Molecular Probes, ThermoFisher Scientific) to label cellular nuclei. Images of the stained sections were obtained with a Leica SP8 laser-scanning confocal microscope equipped with a diode laser 488 nm and 552 nm source for fluorescence illumination with a X-, Y-, and Z-movement controller, and a high-resolution PMT (Zeiss, Oberkochen, Germany) and HyD (Leica) detectors.

### Ex vivo stereological analysis of Cerebral CD13-positive pericyte induced microvessel constrictions and Quantification

Stereological studies were carried out in brain sections to evaluate and quantify the CD13-positive pericyte mediated constrictions in microvessels objectively (19). 10 out of 100 coronal brain sections with a thickness of 50 μm, which were taken 500 μm apart spanning the MCA territory (from the first 1 mm to 6 mm, 5 mm anteroposteriorly in total), were obtained from each brain. To span the MCA territory with unbiased way, coronal sections were obtained with sliding microtome cutting thickness (50 μm) through 5 mm anteroposteriorly starting from 1st mm to 6th. Thus, 1/10 of 100 sections were taken, and in each brain section, for each hemisphere, 10 dissectors with 3D-disector frame (field of view: 240×160 μm along Z-axis (40 μm)) were examined yielding 3.84 mm^2^ as the total area counted. CD13-positive pericyte constrictions were counted through 3D-dissectors within 40 μm sampled thickness. Vessels, pericytes and nuclei were marked with Lectin, anti-CD13, and Hoecsht 33258 (Molecular Probes, ThermoFisher Scientific), respectively. We quantified the number of CD13-positive pericyte induced microvessel constrictions under 40x magnification, defined as a focal narrowing which is more than 20% to diameter of the upstream and/or downstream vessel segment, analyzed whether these constrictions colocalized with pericytes (<10 μm away from the pericyte soma). Microvascular constrictions were counted with among the capillaries having their diameters less than 9 μm and if constrictions were closer than a respective distance for a pericyte soma (<10 μm) were not quantified. The constriction counts were expressed in constrictions/mm^2^.

### PAS Histochemistry and Lectin Staining to Assess Peri-microvascular Glycogen levels

In order to compute the peri-microvascular glycogen, brain sections were labeled for glycogen using periodic acid-Schiff (PAS) staining (11) and vessels were marked by Lectin. 50 μm thick brain sections were permeabilized with Tris buffered saline (TBS) containing 0.3% triton-X for 30 minutes at room temperature, then kept in a solution of 10 mM sodium citrate (pH: 6.0) for 30 minutes heated to 80°C for antigen retrieval. Sections were oxidized in Periodic acid solution (0.5% in double deionized water (ddH20), pH: 7.4) for 10 minutes at room temperature. Since the Schiff’s agent also reacts with non-glycogen aldehyde groups, the sections were kept in dimedone solution (pH: 7.4) at 60°C for 20 minutes. Sections were then incubated with Schiff’s agent for 15 minutes at room temperature and PAS reaction fixed by heated ddH2O for 5 minutes. Lastly, sections were incubated with TBS containing Lectin at + 4°C overnight. Sections were imaged by a fluorescent microscope (400x, Eclipse E600, Nikon Instruments Inc., Melville, NY) equipped with a manually controlled specimen stage for X, Y, and Z-axis, a color camera (model DXM1200, Nikon Instruments Inc.), fluorescent light source (HB-10104AF, Nikon Instruments Inc.), and an image analysis software (NIS-Elements, Version 3.22, Nikon Instruments Inc.). 10 brain sections for each experiment were analyzed using the dissector technique, taking 4-6 dissectors per section (360 × 240 μm) covering the MCA territory (starting 1 mm from the frontal pole to 6 mm of the brain, totally 5 mm anteroposteriorly). A semiautomatic computer routine was used to identify microvessels in the fluorescent channel, and then in the bright field channel quantification of glycogen levels by mean brightness over the selected microvessels were done as established before (19). To make comparisons between the sections possible, each peri-microvascular mean bright field intensity was proportioned to the average background brightfield intensity. Then, correlations among peri-microvascular glycogen levels and total number of microvascular constrictions were analyzed. For this analysis, amount of glycogen around microvessel walls in either DAB injected, or ischemic mice were proportioned to controls. Mean brightness intensity values were assigned on a semi-quantitatively scale (19) ranging 0 to 6 in increments of 1 in DAB injected ones, 0 to 1 in increments of 0.2 in ischemic mice.

### Quantification of Adjusted ischemic infarct volume

Coronal brain sections processed with Nissl staining were evaluated under phase contrast microscope. The ischemic area was delineated and measured by using NIS Elements AR v4.2 software. Adjusted infarct area was calculated by subtracting the non-infarcted area of ipsilateral hemisphere from contralateral hemisphere, and infarct volume was calculated by multiplying the sum of infarct areas of sequential coronal sections of 500 μm apart by total anteroposterior distance of 5 mm (33).

### Statistical analysis

Data were analyzed using IBM SPSS 23 statistical analysis program. All summary data are expressed as mean ± SEM. All data sets were tested for normality using the Shapiro-Wilk normality test. The groups containing normally distributed data were tested using a two-way Student’s t test or analysis of variance (ANOVA). The remaining non-normally distributed data were analyzed using the Mann-Whitney U test (for two groups), Kruskal Wallis (for more than two groups) or “Wilcoxon signed rank” test (for two dependent groups). The Kruskal Wallis and Mann-Whitney U tests were used to compare microvessel constriction counts, and total peri-microvascular glycogen-related brightfield intensity, rCBF changes, ischemic volumes and systolic blood pressure measurements. Wilcoxon signed rank test was used for assessment of systolic blood pressure values within each experiment. Similar variance was assured for all groups, which were statistically compared. Differences with a p < 0.05 were considered to be statistically significant. p values were adjusted for comparison of multiple comparisons using Bonferroni correction.

## RESULTS

### Disruption of CNS glycogen utilization causes CD13-positive pericyte mediated microvascular constrictions in the mouse brain

To investigate the disruption of glycogen utilization pharmacological and genetic methods were used and resultant microvascular impairment was assessed. To show this impairment, CD13-positive pericyte constrictions in the brain microvessels were counted by unbiased semi-stereological quantification parameters as established before in our lab (Supp. Fig. 1A-B) (19).

For pharmacological approach, a potent glycogen phosphorylase inhibitor 1,4-Dideoxy-1,4-imino-D-Arabinitol (DAB) was administered intracerebroventricularly in naïve wild type mice. The effective DAB dose (0.25 M) was decided based on a previous study (25). As the half-life of DAB was found to be 6 hours in *in vitro* enzyme inhibition studies (34), and there were limited data regarding its *in vivo* effects, several time points were examined after i.c.v. DAB injection. To define the possible effects that might be caused by i.c.v. injection or tissue processing, another group of mice received i.c.v. sterile saline (the dissolvent of DAB) as vehicle (n=3).

Mice treated with i.c.v. DAB were sacrificed after 30-min, 1-hr, 3-hr, 6-hr, 9-hr, and 24-hr (n = 3 for all time points). The constrictive effect of DAB injection, counted as mean number of microvascular constrictions (number/mm^2^) were significantly higher compared to i.c.v. vehicle injections at 30-min to 6-hr (p<0.05, Mann-Whitney U test) (Fig. 1 A-B). However, this increase in constrictions within 30min - 6h after administration diminished at 9-h and 24-h, probably due to the diminished effect of DAB which showed that it was not permanent (Fig. 1B). Thus, it was demonstrated that *in vivo* pharmacological inhibition of glycogen phosphorylase resulted in a significant increase in CD13-positive pericyte constrictions of brain microvessels in a probably reversible and time dependent manner.

**Fig.1.**
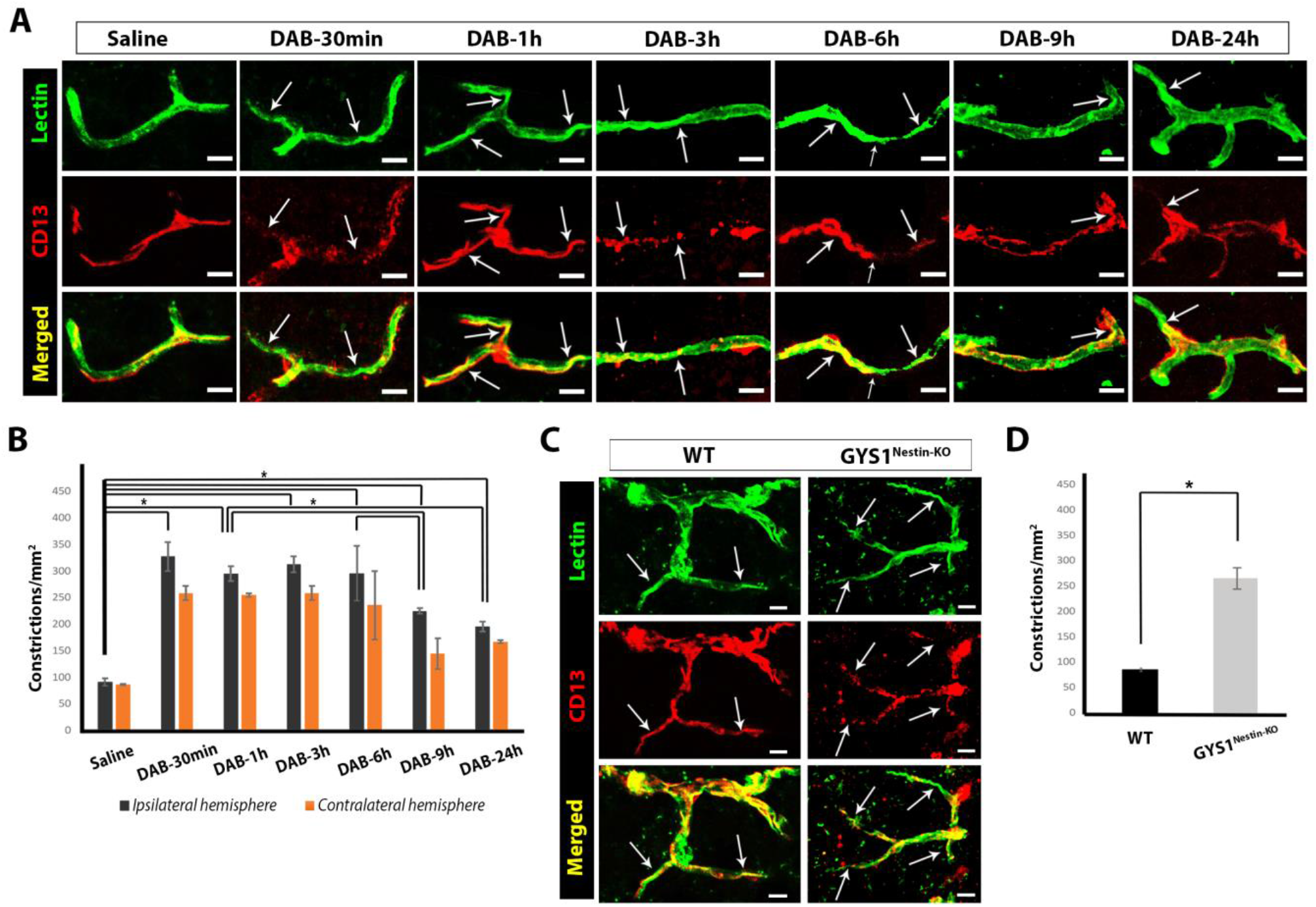
Disruption of CNS glycogen utilization results in CD13 positive (CD13+) capillary pericyte mediated constrictions. CD13 positive capillary pericyte mediated constrictions were evaluated in Swiss albino, wild-type (WT) and GYS1^Nestin-KO^ mice. (A) Experiments performed in adult Swiss albino male and female mice intracerebroventricularly (i.c.v) administered with saline (vehicle) or DAB (as indicated in figure) sacrificed after 30 mins, 1h, 3h, 6h, 9h and 24h (shown left to right respectively). Lycopersicon Esculentum Lectin (upper panel-green) and CD13 (middle panel-red) double labelling (merged as lower panel) reveals increased microvascular constrictions 30 minutes after DAB injections shown by arrows. Images represent 3D reconstruction of 40-μm z-stack. Scale bars, 10 μm. (B) Microvascular constrictions in ipsilateral and contralateral hemispheres were quantified semi-stereologically. DAB (i.c.v) injections result in robust increase of constrictions after 30 minutes, persists for 6 hours. Although number of constrictions started to decline after 9 hours, impact of DAB on these constrictions are not fully reversible even after 24 hours. Quantification of CD13+ pericyte mediated microvascular constrictions in ten fields fields for each hemisphere spanning MCA territory (240 μm × 160 μm) per animal (n = 3 animals per group). Two-way analysis of Kruskal-Wallis and Mann-Whitney U, n = 3; *P < 0.05. Black: ipsilateral hemisphere, Orange: Contralateral hemisphere. (C) Adult naïve wild-type (WT) and GYS1^Nestin-KO^ mice are sacrificed, and brain sections are labeled with Lycopersicon Esculentum Lectin (upper panel-green) and CD13 (middle panel-red and merged as lower panel). GYS1^Nestin-KO^ mice demonstrate higher number of microvascular constrictions as shown by arrows. Images represent 3D reconstruction of 40-μm z-stack. Scale bars, 10 μm. (D) CD13+ pericyte mediated microvascular constrictions in adult naïve wild-type (WT) and GYS1^Nestin-KO^ mice are quantified. Transgenic mice have higher number of constrictions. Quantification of CD13+ pericyte mediated microvascular constrictions in ten fields for each hemisphere spanning MCA territory (240 μm × 160 μm) per animal (n = 3 animals per group). Mann-Whitney U, n = 3; *P < 0.05.

Microvascular constrictions were similar in the neighboring injection areas (1.5-2 mm anteroposterior) and were not different from the corresponding contralateral location (n = 3, p = 0.30, Wilcoxon signed rank test). Moreover, there was no significant difference in the mean number of CD13-positive pericyte mediated microvascular constrictions (number/mm^2^) of ipsilateral compared to contralateral hemispheres within animals of i.c.v. DAB (p=0.109) and vehicle (p=0.109, Wilcoxon signed rank test) for all time points. These results suggested that DAB distributed all over the brain from the i.c.v. injection site and was effective.

Next, to determine the direct effect of peri-microvascular glycogen we used GYS-1^Nestin-KO^ mice. In GYS-1^Nestin-KO^ mice, the enzyme glycogen synthase-1 responsible for the production of glycogen, was selectively deleted from the brain (26). CD13-positive pericyte mediated microvessel constrictions (number/mm^2^) of naïve GYS-1^Nestin-KO^ mice displayed significantly higher microvascular constrictions (260.85 ± 20.87, n=3) when compared to wild type naïve mice (88.72 ± 3.00, n=3) (p=0.025, Mann-Whitney U test) (Fig. 1C-D).

### Glycogen Phosphorylase Inhibition Disrupts Glycogen Utilization Where CD13-positive pericyte mediated Microvascular Constrictions Occur

Periodic acid Schiff (PAS) staining was used as it is the gold standard to show glycogen in the tissue. Using an aldehyde blocker dimedone, allowed us to visualize only glycogen, but not the glycoproteins and proteoglycans present in the tissue (35). Thereafter, a semi-automatic computer software (’Macro’, NIS Elements 4.3) that was previously established in our laboratory was used to quantify the peri-microvascular glycogen (19). In this method, the peri-microvascular brightfield signal intensity was related to the glycogen content. DAB treated brains (n = 3/group) displayed a significant increase in the mean brightfield intensity of PAS signal (Fig. 2A-B) within 1 hr (10.28 ± 1.11), 6-hr (8.82 ± 1.54), and 24-hr (6.13 ± 0.51) compared to vehicle treated group (2.47 ± 0.32, p= 0.025 for each, Mann-Whitney U test) (Fig. 2C-D). The increase coincided the location of microvascular constrictions, hence glycogen is present there, but possibly because of the disrupted glycogen utilization in the peri-microvascular astrocyte processes, cannot be utilized by glycogen phosphorylase.

**Fig. 2.**
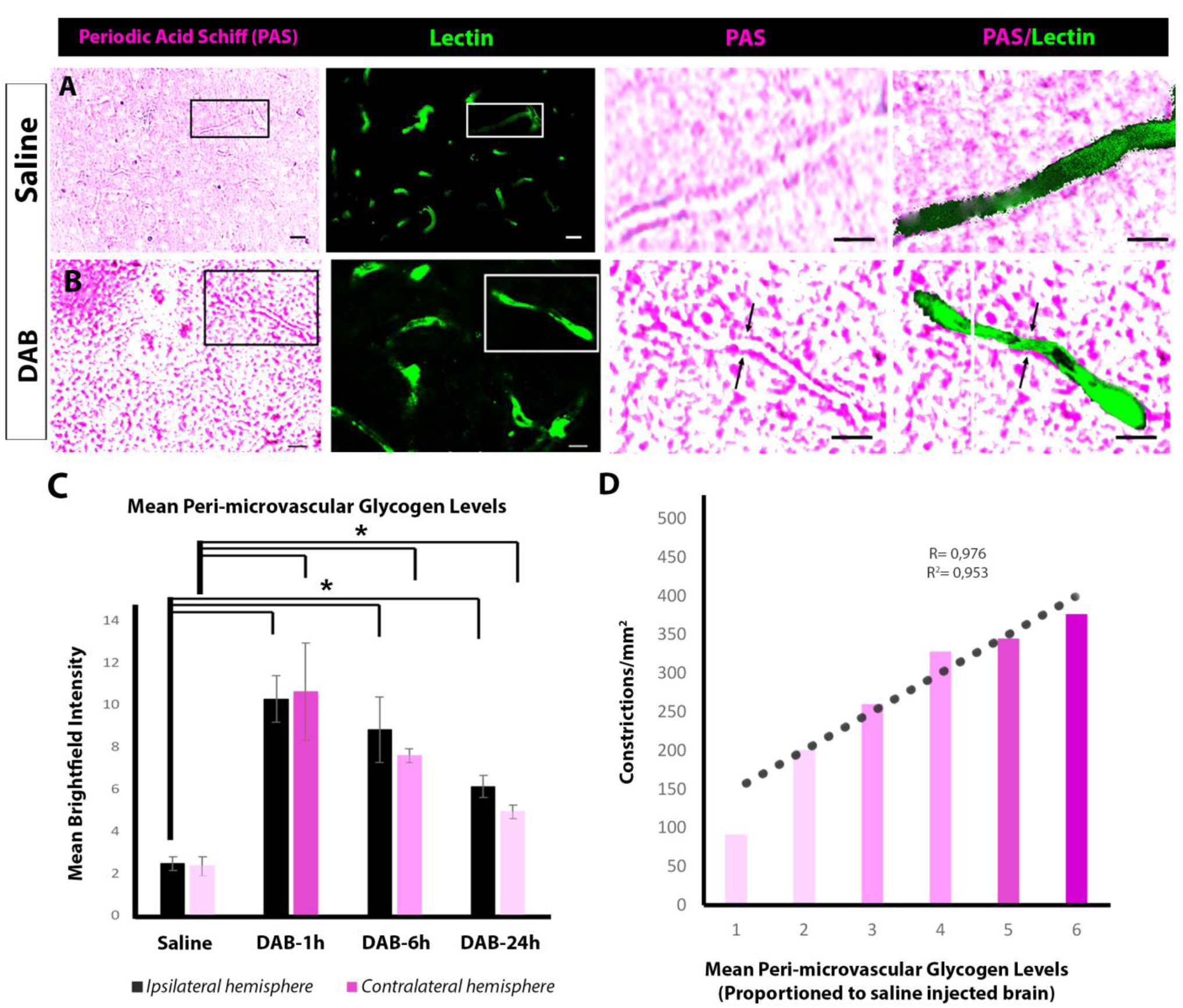
Intracerebroventricularly administered glycogen phosphorylase inhibitor 1,4-Dideoxy-1,4-imino-D-arabinitol hydrochloride (DAB) causes increased peri-microvascular glycogen and correlates with microvascular constrictions. Periodic acid Schiff (PAS)-Lycopersicon esculentum Lectin double staining are performed in i.c.v saline or DAB injected Swiss albino mice to show glycogen around microvessels in the tissue. (A) PAS (left panel) and Lectin (second from the left) stained section from saline injected mice reveal stored glycogen around microvessels (black and white box). The areas delineated via boxes are displayed with higher magnification and these images show peri-microvascular glycogen (third and fourth panels from the left). Scale bars; 20 μm (left and second to left), 10 μm (third and fourth to left). (B) DAB injection causes increase of glycogen deposition around microvessels (left panel-black box) coinciding with microvessel constrictions (second from the left-white box). Magnified areas display increased glycogen related brighfield intensity especially around microvascular constrictions (arrows). Scale bars; 20 μm (left and second to left), 10 μm (third and fourth to left). (C) A semi-automatic software macro (NIS Elements 4.3) that was previously established in our laboratory is used to quantify the peri-microvascular glycogen (Alarcon-Martinez et al, Acta Neuropath. Comm. 2019). The peri-microvascular brightfield signal intensity is measured in both hemispheres after injections. DAB treated brains (n = 3 animals per group) displayed a significant increase in the mean brightfield intensity of PAS signal compared to saline (Two-way analysis of Mann-Whitney U, n = 3; *P < 0.05). Black: ipsilateral hemisphere, Magenta: Contralateral hemisphere. (D) The total number of constrictions (y-axis) are positively correlated to the mean peri-microvascular glycogen levels (x-axis). Hence glycogen is there but cannot be utilized by glycogen phosporylase. Pearson’s correlation, R= 0.976, R^2^ = 0.953.

To further investigate the role of glycogen utilization in CD13-positive pericyte mediated microvessel constrictions, correlations among peri-microvascular glycogen levels and total number of CD13-positive pericyte mediated microvascular constrictions were analyzed. For this analysis, amount of glycogen around microvessel walls in DAB injected mice were proportioned to vehicle injected controls. Mean brightness intensity values were assigned on a semi-quantitative manner (19). Interestingly, DAB induced CD13-positive pericyte mediated constrictions tightly correlated with higher glycogen levels (Pearson’s correlation, R= 0.976, R^2^ = 0.953, (Fig. 2D). Therefore, these results suggested that disruption of glycogen utilization via glycogen phosphorylase inhibition exacerbated CD13-positive pericyte mediated microvascular constrictions.

### Cerebral Ischemia Depletes Peri-microvascular Glycogen and Increases DAB Related CD13-positive pericyte mediated Microvascular Constrictions Further

To evaluate the effect of cerebral ischemia on glycogen utilization, 1 hr after i.c.v. vehicle (saline) injection, permanent MCAo was performed for 2-hr (n = 3). A decrease in PAS staining (very light pink color), depicting glycogen reduction, was observed throughout the MCA territory corresponding to the infarction (Figure 3A) (1.11 ± 0.35), and was significantly lower when compared to contralateral hemisphere (2.12 ± 0.42, p = 0.025, Mann-Whitney U test), as well as non-ischemic vehicle injections (2.47 ± 0.32, n = 3, p = 0.025). In mice subjected to 2-hr permanent MCAo after DAB injection (n=3), we determined an increase in peri-microvascular glycogen (Fig.ure 3B) (4.10 ± 1.02) compared to both non-ischemic vehicle (p = 0.025), and ischemic vehicle injected groups (1.11 ± 0.35, p = 0.025, Mann-Whitney U test) in both hemispheres (p = 0.011, Kruskal-Wallis test) (Fig. 3C).

**Fig. 3.**
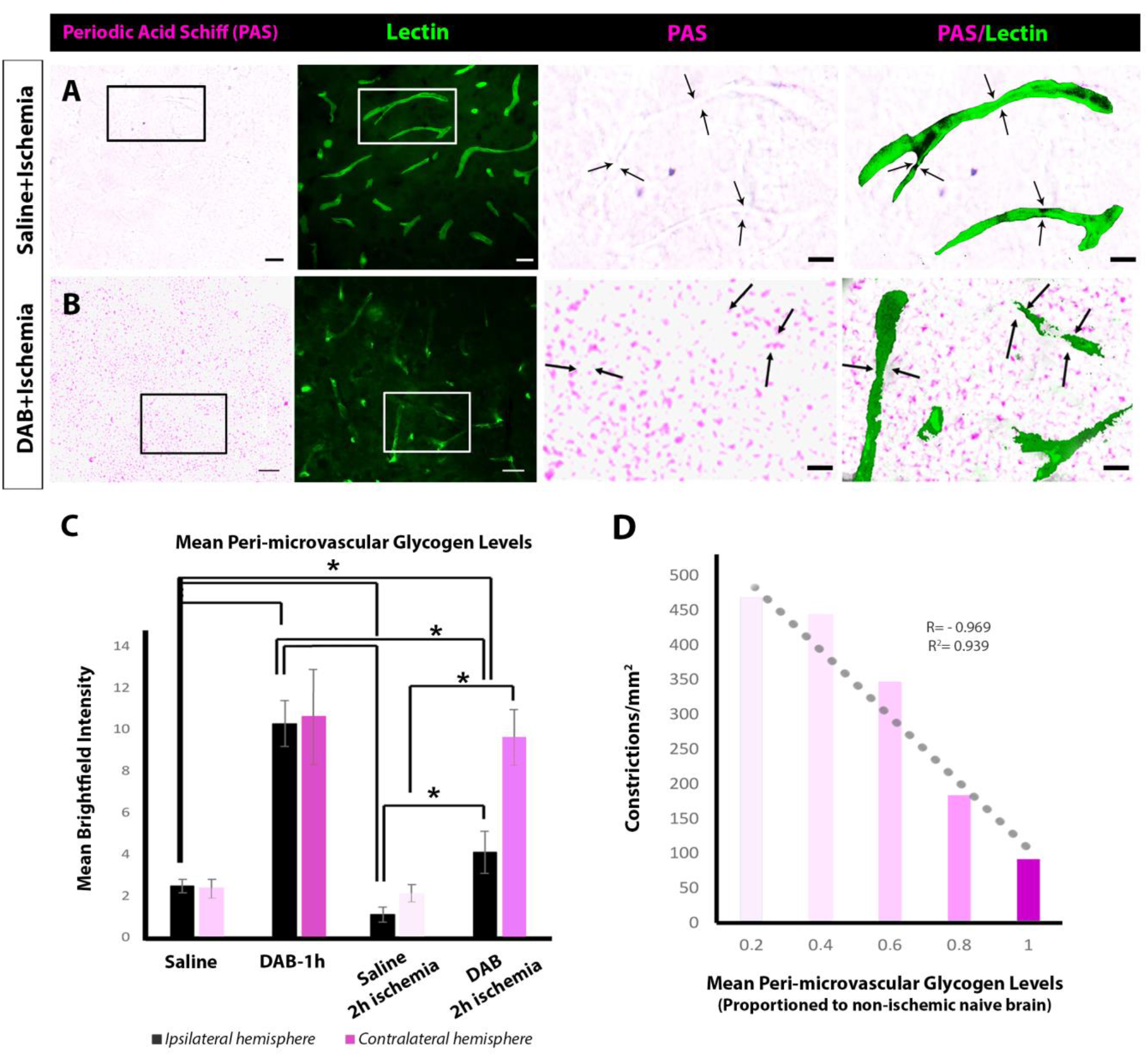
Cerebral Ischemia Depletes Peri-microvascular Glycogen and Increases Microvascular Constrictions after glycogen phosphorylase inhibition. Periodic acid Schiff (PAS)-Lycopersicon esculentum Lectin double staining are performed in Swiss albino mice which underwent MCAo after i.c.v saline or DAB injections (n = 3 animals per group). (A) PAS (left panel) and Lectin (second from the left) stained section from saline injected ischemic mice reveal depletion of stored glycogen around microvessels (black and white box). The areas delineated via boxes are displayed with higher magnification and these images show diminished peri-microvascular glycogen (third and fourth panels from the left) coinciding the constrictions (arrows). Scale bars; 20 μm (left and second to left), 10 μm (third and fourth to left). (B) In mice subjected to 2-hr permanent MCAo after DAB injection (n=3), there is an increase in perimicrovascular glycogen (left panel-black box) coinciding with higher number of microvessel constrictions (second from the left-white box). Magnified areas display increased glycogen related brighfield intensity especially around microvascular constrictions (arrows). Scale bars; 20 μm (left and second to left), 10 μm (third and fourth to left). (C) Quantification of mean peri-microvascular glycogen levels via a semi-automatic software macro (NIS Elements 4.3) is used (Alarcon-Martinez et al, Acta Neuropath. Comm. 2019). Mean brightfield intensity of DAB treated brains (n = 3 animals per group) displayed a significant increase in the mean brightfield intensity of PAS signal compared to saline (Two-way analysis of Mann-Whitney U, n = 3; *P < 0.05). Black: ipsilateral hemisphere, Magenta: Contralateral hemisphere. (D) The total number of constrictions (y-axis) are correlated to the decrease of mean peri-microvascular glycogen levels (x-axis). Although DAB blocked the utilization of glycogen, ischemia depletes and causes higher number of microvascular constrictions. Pearson’s correlation, R= −0.969, R^2^ = 0.939.

Correlations among peri-microvascular glycogen levels and total number of CD13-positive pericyte mediated microvascular constrictions were assessed in ischemic and non-ischemic controls. Values were assigned on a semi-quantitative manner (19). After 2-h of permanent ischemia, the number of CD13-positive pericyte mediated constricted microvessels correlated with the presence of low perimicrovascular glycogen levels (Pearson’s correlation, R= −0.969, R^2^ = 0.939, (Fig. 3D). Therefore, these results also suggest that depletion of glycogen due to ischemia results in CD13-positive pericyte mediated microvascular constrictions and aggravated more by the inhibition of glycogen phosphorylase.

In addition to PAS histochemical staining, immunohistochemical studies were performed to show stored glycogen by using non-commercial ESG1A9 and IV58B6 antibodies that label glycogen granules, thus glycogen stores. Glycogen is mostly stored in peri-microvascular astrocyte processes (36). Glycogen labelling was evident in the astrocyte processes covering the microvascular walls, where constrictions were not present (Supp. Fig. 2B). Following 2-hr permanent MCAo, there was a drastic reduction of peri-microvascular glycogen labelling in the ischemia core area (Supp. Fig. 2D), and an increase in the peri-infarct area, which was similar to PAS histochemistry (Supp. Fig. 2E). Moreover, glycogen labelling was absent in the ischemia-induced microvessel constriction locations (Supp. Fig. 2D-E). When DAB is administered, peri-microvascular glycogen labelling drastically increased (Supp. Fig. 2F-G) and coincided with the microvascular constrictions.

### Glycogen utilization affects cerebral blood flow

To investigate the functional consequences of the relationship between peri-microvascular glycogen and CD13-positive pericyte mediated microvascular constrictions in the brain, we studied regional cortical cerebral blood flow changes under physiological and pathological conditions (Fig. 4). Regional cortical blood flow (rCBF) alteration maps were created by the semi-automatic analysis routine (MATLAB-Simulink) with the data obtained by Laser Speckle contrast imaging during MCA occlusion (Fig. 4A-B). Laser Speckle contrast imaging was performed throughout ipsilateral permanent MCAo before, during and 1 hr after the occlusion.

**Fig.4.**
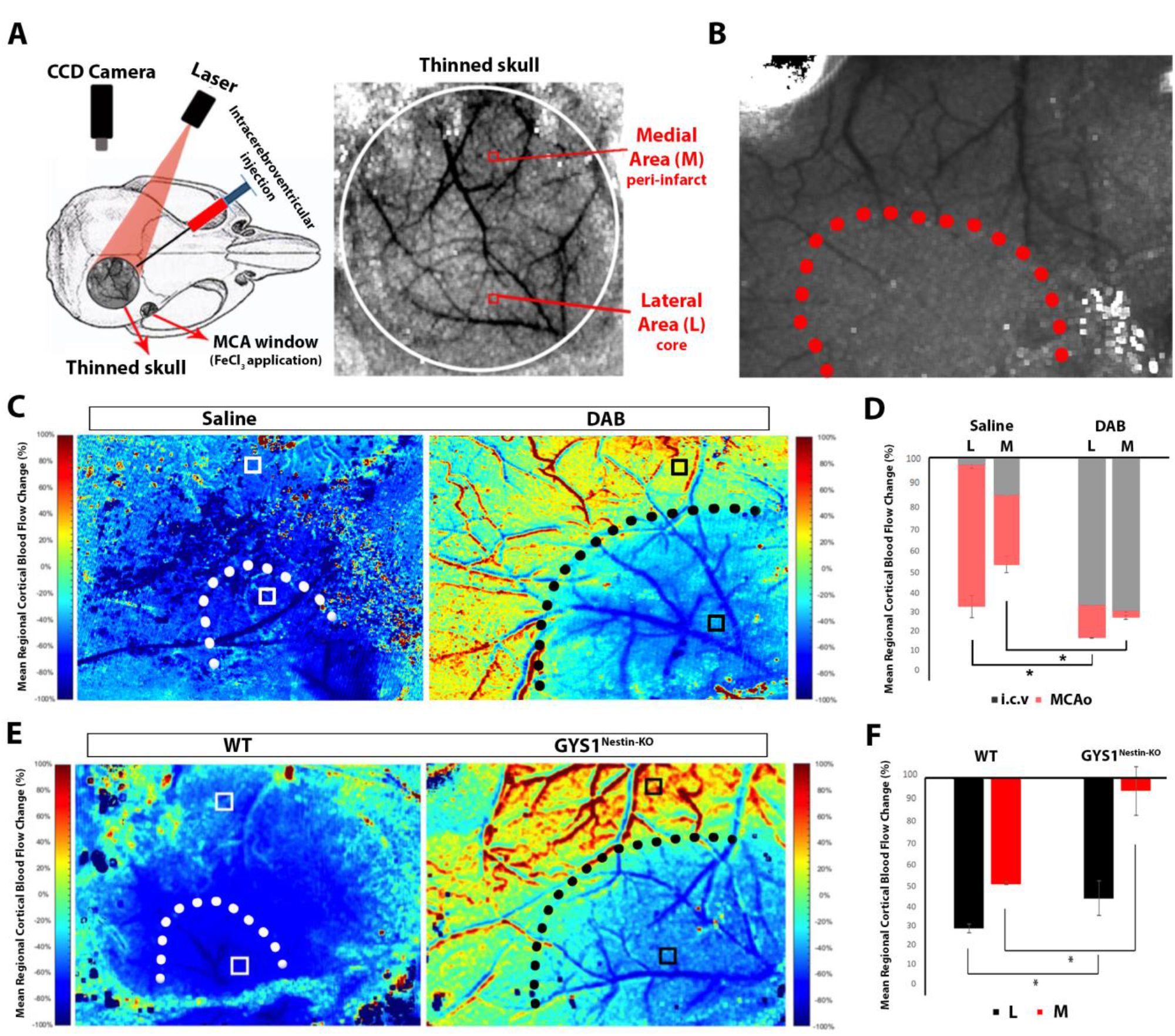
Glycogen utilization affects cortical regional Cerebral Blood Flow (rCBF) dynamics after successful Middle Cerebral Artery occlusion (MCAo). Laser speckle contrast imaging (LSCI) is used to detect the cortical blood flow changes observed through the cranial window as described previously (Dunn et al 2001). (A) Schematic illustration of experimental setup (left panel). DAB and saline injections via i.c.v are also performed through cranial window. MCAo is introduced through another cranial window via FeCl3 administration for 10 minutes. CCD camera captures the reflection of laser illuminated cortical region through thinned skull and then rCBF changes observed throughout the whole experiments as described. The visualization of cortical surface is demonstrated on the right panel. Vessels appear as black, whereas parenchyme is gray. The medial (depicting peri-infarct area) and lateral (ischemic core) region of interests (ROI) are carefully chosen to avoid high caliber vasculature. (B) Representative image taken during MCAo. Red dots delineate the area of ischemic core after successful clotting of MCA. (C) Pseudocolored maps are created after MCAo of saline (left panel) and DAB (right panel) injected WT mice. Every LSCI image taken 10 seconds apart proportioned to baseline after i.c.v injections (described in Methods). Scale bars on the right and left sides show that cold colors represent a decrease (0 through -%100), whereas hot colors, an increase in rCBF from baseline (0 through +%100). The quantifications of mean rCBF change are perfomed in white (saline) and black (DAB) boxes from medial (peri-infarct) and lateral (core) ROIs. White (saline) and black (DAB) dots describe the MCA core territories, respectively. Drift in z axis (vessel shift) was inevitable because of the blood volume changes in the ischemic tissue. (D) Quantifications of mean rCBF change in each ROI. Results are given as percentages (n = 3 animals per group). The average values of mean rCBF changes measured at medial (M) and lateral (L) regions of interest (white and black boxes) are shown as gray after i.c.v saline or DAB injections, red after MCAo. DAB injection resulted a robust reduction of mean rCBF compared to saline within an hour (Two-way analysis of Mann-Whitney U, n = 3; *P < 0.05). Ipsilateral MCAo for 1 hr, decreased mean rCBF further in the ischemic core of DAB injected mice: however, lower than vehicle injected controls (Two-way analysis of Mann-Whitney U, n = 3; *P < 0.05). In DAB injected ischemic mice, peri-infarct ROI displays less reduction of mean rCBF compared to vehicle injected ischemic controls (Two-way analysis of Mann-Whitney U, n = 3; *P < 0.05). (E) rCBF alterations are also assessed in GYS1^Nestin-KO^ mice (right panel) during permanent MCAo compared to WT (left panel) via pseudocolored maps. The representations are same as above. GYS1^Nestin-KO^ mice display similar rCBF alteration pattern (right panel) as DAB injected counterparts (right panel at C). (F) Quantifications of mean rCBF change in each ROI. Results are given as percentages (n = 3 animals per group). The average values of mean rCBF changes measured at medial (M) and lateral (L) regions of interest (white and black boxes) are shown as red and black after MCAo, respectively. Ischemia caused a decrease in rCBF in the ischemic core of both WT and GYS1^Nestin-KO^ mice, but the decrease was less in GYS1^Nestin-KO^ group (Two-way analysis of Mann-Whitney U, n = 3; *P < 0.05). Besides, the mean decrease was also significantly lower in peri-infarct area in wild type mice, compared to GYS1^Nestin-KO^ (Two-way analysis of Kruskal-Wallis and Mann-Whitney U, n = 3; *P < 0.05).

DAB injection resulted a robust reduction of mean rCBF (medial: 71.54 ± 0.58 %, lateral: 68.81 ± 0.29 %) compared to vehicle injection (medial: 17.07 ± 0.13%, lateral: 2.37 ± 1.86%, n=3, p= 0.025, Mann-Whitney U test) within an hour (Fig. 4C). Ipsilateral MCAo for 1 hr, decreased mean rCBF further in the ischemic core of DAB injected mice (50.23 ± 1.03%); however, lower than vehicle injected controls (69.22 ± 5.36%, p= 0.025, Mann-Whitney U test) (Fig. 4D). In DAB injected ischemic mice, peri-infarct area displayed less reduction of mean rCBF (11.66 ± 3.71%) compared to vehicle injected ischemic controls (39.45 ± 4.57%, p= 0.025, Mann-Whitney U test) (Fig. 4D) possibly because of the maximum detectable reduction of rCBF after DAB.

rCBF alterations were assessed in GYS-1^Nestin-KO^ mice during ipsilateral permanent MCAo compared to wild type (C57Bl/6J) controls via laser speckle contrast imaging as well (Fig. 4E-F). Ischemia caused a decrease in rCBF in the ischemic core of both wild type (70.92 ± 2.10%, n=3) and GYS-1^Nestin-KO^ (56.76 ± 8.09%, n=3) mice, but the decrease was less in GYS-1^Nestin-KO^ group (p = 0.025, Mann-Whitney U test) (Fig. 4E-F). Besides, the mean decrease was also significantly lower in periinfarct area in wild type mice (49.83 ± 0.71%), compared to GYS-1^Nestin-KO^ (6.19 ± 1.58%, p = 0.025, Mann-Whitney U test) (Fig. 4E-F) in accordance with DAB and ischemia results. These results suggest that GYS-1^Nestin-KO^ mice may have a less basal rCBF level resulting less rCBF decrease after ischemia.

As the physiological parameters of mice have a notable effect on the outcome measures such as hemodynamic differences and infarct volume, it was highly critical to standardize parameters like systolic blood pressure in the peri-experimental setting, hence it was closely monitored (Table 1). Systolic blood pressure was measured non-invasively in the experiments by LSC imaging, using a tail cuff, in order not to cause a probable hemodynamic change due to inevitable blood loss in conventional invasive measurements. No significant changes were observed between the mean systolic blood pressure values measured before and after the permanent MCAo among groups (Table 1).

### Disrupted peri-microvascular glycogen increases susceptibility to ischemia

Ischemic infarct volumes after MCAo were significantly higher in DAB treated mice (18.61 ± 2.22) than vehicle treated groups (10.72 ± 0.42 mm^3^) (n =3/groups) (p =0.025, Mann-Whitney U test). Infarct volumes of GYS-1^Nestin-KO^ group were also higher (21.22 ± 1.52) when compared to wild type (8.58 ± 0.05 mm^3^)(n =3/groups) (p = 0.025, Mann-Whitney U test) (Fig. 5A-B). This increase in infarct volume further pointed out the vulnerability of mice with disrupted peri-microvascular glycogen to ischemia. Accordingly, DAB injected mice (462.11 ± 8.40/mm^2^) and GYS-1^Nestin-KO^ (430.45± 8.13/mm^2^) had prevalent CD13-positive pericyte mediated microvascular constrictions in the ischemic area (Fig. 5C-D) compared to vehicle injected (371.85 ± 32.42) and wild type mice (384.28 ± 30.16, n=3/groups, p= 0.025, Mann-Whitney U test) (Fig. 5C-D). Ischemia further resulted in higher number of CD13-positive pericyte mediated microvascular constrictions in mice with disrupted glycogen (DAB injected and GYS-1^Nestin-KO^) compared to ischemic controls (p= 0.001, Kruskal-Wallis test). Therefore, disruption of glycogen utilization increased further susceptibility during ischemia.

**Fig.5.**
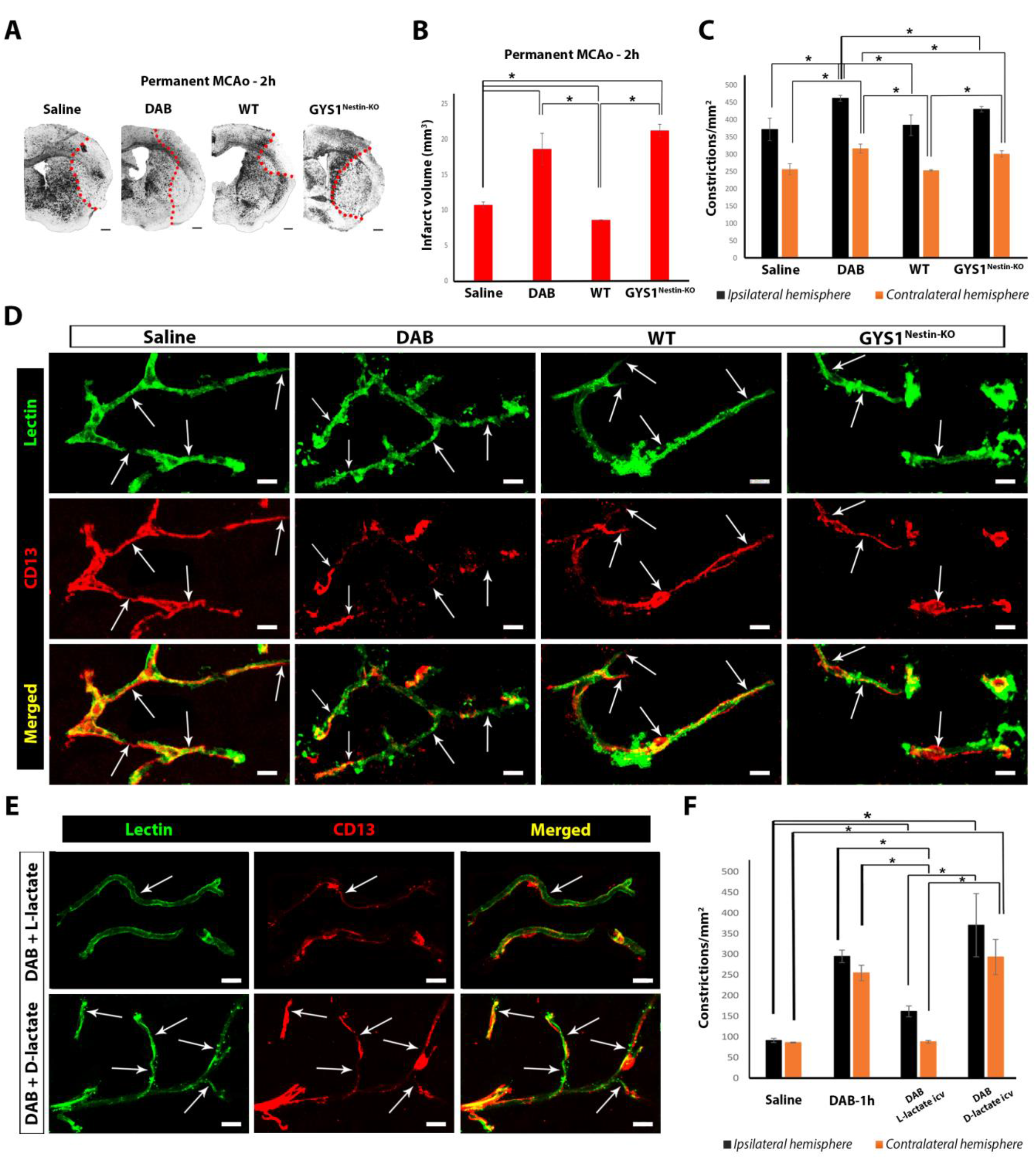
Susceptibility to cerebral ischemia is potentiated via disruption of CNS glycogen utilization which is reversible. Permanent MCAo experiments performed in Saline, DAB injected WT, naïve WT and GYS1^Nestin-KO^ mice (n = 3 animals per group). (A) Representative images taken from Cresyl-violet stained sections of ischemic mice via phase contrast microscopy (as indicated in figure) that underwent 2-hour MCAo. Red dots are placed over the border between core and peri-infarct areas. Scale bars, 500 μm. (B) Infarct volumes after 2-hour MCAo are measured then quantified with volume correction (Swanson, JCBFM 1990). Ischemic infarct volumes after MCAo were significantly higher in DAB treated mice than saline treated groups (Two-way analysis of Mann-Whitney U, n = 3; *P < 0.05). Infarct volumes of GYS1^Nestin-KO^ group were also higher when compared to wild type (Two-way analysis of Mann-Whitney U, n = 3; *P < 0.05). (C) Quantification of microvascular constrictions after MCAo in Saline, DAB injected WT, naïve WT and GYS1^Nestin-KO^ mice (n = 3 animals per group). Quantification of CD13+ pericyte mediated microvascular constrictions are held in ten fields fields for each hemisphere spanning MCA territory (240 μm × 160 μm) per animal. Ischemia further resulted in higher number of microvascular constrictions in peri-microvascular glycogen disrupted mice (DAB injected and GYS1^Nestin-KO^) compared to ischemic controls (Two-way analysis of Mann-Whitney U, n = 3; *P < 0.05). Black: ipsilateral hemisphere, Orange: Contralateral hemisphere. (D) Lycopersicon Esculentum Lectin (upper panel-green) and CD13 (middle panel-red) double labelling (merged as lower panel) reveals increased microvascular constrictions (arrows) 2 hours after MCAo in Saline, DAB injected WT, naïve WT and GYS1^Nestin-KO^ mice (left to right, respectively). Images represent 3D reconstruction of 40-μm z-stack. Scale bars, 10 μm. (E) Lycopersicon Esculentum Lectin (left panel-green) and CD13 (middle panel-red) double labelling (merged as right panel) in i.c.v DAB+ L/D-lactate injected mice. Double labelling shows that L-lactate (upper panel) reverses the DAB’s impact on CD13+ pericyte mediated constrictions contrary to its enantiomer D-Lactate (lower panel). Images represent 3D reconstruction of 40-μm z-stack. Scale bars, 10 μm. (F) Quantification of CD13+ pericyte mediated microvascular constrictions in i.c.v DAB+ L/D-lactate injected mice compared to controls (n = 3 animals per group). L-lactate can reverse the DAB-induced constrictions; however, D-lactate demonstrates similar number of microvascular constrictions with DAB (Two-way analysis of Mann-Whitney U, n = 3; *P < 0.05). Black: ipsilateral hemisphere, Brown: Contralateral hemisphere.

### L-lactate administration reversed DAB-induced CD13-positive pericyte mediated microvascular constrictions

Next, we tested whether CD13-positive pericyte mediated constrictions after DAB are reversible. Energy metabolite transferred from astrocytes to neurons upon glycogen breakdown is lactate (2). Therefore, we used lactate as an alternative energy source to replace glycogen (25). L-lactate and its optical isomer D-lactate were administered intracerebroventricularly. L-lactate was able to reverse the DAB-induced CD13-positive pericyte mediated constrictions (ipsilateral: 161.89/mm^2^ ± 13.28, contralateral: 81.16 ± 1.25/mm^2^); however, D-lactate (ipsilateral: 370.66 ± 76.68/mm^2^, contralateral: 293.40 ± 42.60/mm^2^) (Fig. 5E-F) demonstrated similar number of CD13-positive pericyte mediated microvascular constrictions with DAB (n=3, p= 0.025, Mann-Whitney U test).

## Discussion

The mechanisms involved in the microvascular constrictions, which adversely affect tissue perfusion, have not yet been fully elucidated. We found that disruption of glycogen mobilization either pharmacologically or genetically, via DAB and GYS-1^NestinKO^ respectively lead to microvascular constrictions, ultimately suggesting that disturbed glycogen metabolism can cause ischemic-like phenotype in the brain under non-ischemic circumstances. Although it has been shown in various studies that glycogen has essential roles in the brain, investigating the regional differences has been an issue because of the difficulties of quantification, and demonstrating in the tissue (37–39). Our ex vivo data which was analyzed by semi-stereological methods to overcome this regional heterogeneity showed that CD13-positive pericytes were localized at constriction sites in accordance with our previous results showing that pericytes have an important role causing microvascular constrictions after cerebral and retinal ischemia (17, 19). After brain ischemia, the extracellular potassium increases which preceded by prompt Ca^2+^ flux and a decrease in ATP, eventually inducing the activation of phosphorylase kinase (40). Therefore, brain glycogen is degraded to meet this sudden energy demand when the rate of glycogenolysis may rise approximately 200-fold above that in resting state (3, 41, 42). This might explain why inhibiting glycogenolysis or abolishing glycogenesis mimicked brain ischemia. Interestingly, it was also demonstrated that DAB induced CD13-positive pericyte mediated constrictions in brain with a reversible and time dependent manner.

Glycogen utilization was inhibited with DAB and excess glycogen was detected both with PAS and non-commercial, glycogen-targeted specific antibodies (ESG1A9 and IV58B6) immunofluorescently around the close vicinity of CD13-positive pericyte mediated microvascular constrictions (35, 39). Preventing utilization of energy metabolites from glycogen with DAB, there was an increase in glycogen related signals despite increased number of CD13-positive pericyte mediated constrictions. This suggests that DAB caused increased glycogen content at the astrocyte end-feet and around the CD13-positive pericyte mediated microvascular constrictions by inhibiting glycogen phosphorylase. This might be an explanation how glycogen increases in the periinfarct area in accordance to a recent study linking the relationship between increased glycogen in the periinfarct area of human, monkey and mouse brain and glycogen phosphorylase activation via insulin mediated mechanism in cerebral ischemia reperfusion model (43).

We also found that there was a decrease in the amount of glycogen around microvessels shown especially where constrictions took place in ischemic condition similar to constrictions in similar to those reported for ischemic retinal microvessels (19). As a result of 2 hours of ischemia, glycogen around the microvascular niche was rapidly utilized in the infarct core and caused CD13-positive pericyte mediated constrictions in accordance with previous results showing PAS-stained tissue sections revealed a lack of glycogen within the ischemic infarct (27). The constriction of microvessels induced by ischemia were shown to be correlated with reduction of perivascular glycogen in retinal glycogen, showing that early phases of ischemic stroke, glycogen utilization plays an important role in functional regulation of neurogliovascular unit in microcirculatory disorders (19, 43). Of note, the amount of glycogen was not quantified in order not to speculate since microwave fixation needs to be performed to preserve glycogen while harvesting the tissues (22, 39, 41, 44).

In addition, DAB injected, and GYS1^Nestin-KO^ mice displayed different cortical cerebral blood flow alteration maps compared to controls after inducing permanent MCAo. Mean regional cortical blood flow (rCBF) changes in both conditions where glycogen metabolism was abolished decreased significantly in the ischemic core compared to controls. Interestingly, the percentages of reduction of overall rCBF in the core areas of both DAB treated and GYS1^Nestin-KO^ mice were much lower than controls. Both because only DAB treatment even caused significant rCBF reduction and core rCBF reduction was significantly less than WT controls, we might speculate that GYS1^Nestin-KO^ mice displayed much lower basal capillary blood flow under normal circumstances. These findings warrant the further cerebral blood flow studies. On the other hand, peri-infarct areas did not show significant decrease in average blood supply as in the controls. This was further supported by an increase in ischemic infarct volume in these mice even after 2-hour of ischemia.

Moreover, the peri-microvascular glycogen utilization in the brain has been shown to significantly aggravated the ischemia-induced persistent CD13-positive pericyte mediated microvascular constrictions. As the glycogen phosphorylase inhibition was sustained, these constrictions took place throughout MCA territory. Thus, the ischemic tolerance of these mice was diminished, and the constrictions occurred higher in cerebral ischemia and hence the infarct volume increased. On the contrary, one previous study showed that inhibiting glycogen phosphorylase with CP-316,819 attenuated ischemic tissue damage and reduced the infarct size (45). This seemingly contradictory results could be addressed that CP-316,819 interestingly cannot exert its inhibitory effect on glycogen phosphorylase under ischemic condition which might have enabled the utilization of increased glycogen content in brain. In other words, CP-316,819 treatment first increased the glycogen content in brain, and this met the energy requirements longer than controls by allowing the cells utilize glycogen when ischemia was induced (12).

Based on the reports showing that lactate derived from glycogen stored at astrocyte processes sustain the support to glutamatergic neuronal synapses named as ‘Astrocyte neuron lactate shuttle (ANLS)’ (39, 40), we hypothesized that the astrocyte end-feet surrounding microvascular wall may also sustain pericytes to keep the capillaries patent especially under ischemic conditions where energy requirements are unmet. Also, astrocytes with intact plasma membranes, hence viable and resistant to hypoxia in the ischemic area was determined to be positive for glycogen (11). Therefore, we employed L-lactate and found that it reversed DAB-induced CD13-positive pericyte mediated microvascular constrictions, suggesting that pericyte constrictions are mediated by the availability of releasable lactate derived from stored glycogen in astrocyte end-feet. Although revealing the underlying mechanisms requires further research, we can speculate that glycogen stores in astrocytes sustain pericyte contractibility thus adequate tissue perfusion especially under demanding conditions. These results were in accordance with the previous study demonstrating that the application of L-lactate reversed the decrease in the cortical spreading depolarisation threshold induced by DAB (25) and reduced infarct size with improved neurological outcomes after MCAo (29).

We addressed that peri-microvascular glycogen plays an important role in the components that can cause a susceptibility for many CNS diseases, primarily in the very early ischemic phase by mediating CD13-positive pericyte mediated microvascular constrictions, thus adequate tissue perfusion. Our study on the pericyte behaviour in relation to glycogen metabolism and cerebral blood flow in health and disease may also give additional insight to the scientists who have been researching on glycogen and its importance in the pathophysiology of many diseases such as ischemic stroke, migraine, cognitive dysfunction, and memory loss (25, 26, 36, 43, 44). These observations specifically focusing on microvascular niche also warrant further studies to understand the metabolic crosstalk between glial end-feet and pericytes in the control of microvascular function.

## Notes

### Competing Interest Statement

The authors have declared no competing interest.

## References

1. Attwell D, Laughlin SB. An energy budget for signaling in the grey matter of the brain. J Cereb Blood Flow Metab. 2001;21(10):1133–45.

2. Magistretti PJ, Allaman I. A cellular perspective on brain energy metabolism and functional imaging. Neuron. 2015;86(4):883–901.

3. Dringen R, Gebhardt R, Hamprecht B. Glycogen in astrocytes: possible function as lactate supply for neighboring cells. Brain Res. 1993;623(2):208–14.

4. Pellerin L, Bouzier-Sore AK, Aubert A, Serres S, Merle M, Costalat R, et al. Activity-dependent regulation of energy metabolism by astrocytes: an update. Glia. 2007;55(12):1251–62.

5. Pellerin L, Magistretti PJ. Glutamate uptake into astrocytes stimulates aerobic glycolysis: a mechanism coupling neuronal activity to glucose utilization. Proc Natl Acad Sci U S A. 1994;91(22):10625–9.

6. De Keyser J, Steen C, Mostert JP, Koch MW. Hypoperfusion of the cerebral white matter in multiple sclerosis: possible mechanisms and pathophysiological significance. J Cereb Blood Flow Metab. 2008;28(10):1645–51.

7. Lipton P. Ischemic cell death in brain neurons. Physiol Rev. 1999;79(4):1431–568.

8. Bouma GJ, Muizelaar JP, Stringer WA, Choi SC, Fatouros P, Young HF. Ultra-early evaluation of regional cerebral blood flow in severely head-injured patients using xenon-enhanced computerized tomography. J Neurosurg. 1992;77(3):360–8.

9. Magistretti PJ, Pellerin L, Rothman DL, Shulman RG. Energy on demand. Science. 1999;283(5401):496–7.

10. Saez I, Duran J, Sinadinos C, Beltran A, Yanes O, Tevy MF, et al. Neurons have an active glycogen metabolism that contributes to tolerance to hypoxia. J Cereb Blood Flow Metab. 2014;34(6):945–55.

11. Gurer G, Gursoy-Ozdemir Y, Erdemli E, Can A, Dalkara T. Astrocytes are more resistant to focal cerebral ischemia than neurons and die by a delayed necrosis. Brain Pathol. 2009;19(4):630–41.

12. Obel LF, Muller MS, Walls AB, Sickmann HM, Bak LK, Waagepetersen HS, et al. Brain glycogen-new perspectives on its metabolic function and regulation at the subcellular level. Front Neuroenergetics. 2012;4:3.

13. Fern R. Ischemic tolerance in pre-myelinated white matter: the role of astrocyte glycogen in brain pathology. J Cereb Blood Flow Metab. 2015;35(6):951–8.

14. Rothman DL, Dienel GA. Development of a Model to Test Whether Glycogenolysis Can Support Astrocytic Energy Demands of Na(+), K(+)-ATPase and Glutamate-Glutamine Cycling, Sparing an Equivalent Amount of Glucose for Neurons. Adv Neurobiol. 2019;23:385–433.

15. Bak LK, Walls AB, Schousboe A, Waagepetersen HS. Astrocytic glycogen metabolism in the healthy and diseased brain. J Biol Chem. 2018;293(19):7108–16.

16. Peppiatt CM, Howarth C, Mobbs P, Attwell D. Bidirectional control of CNS capillary diameter by pericytes. Nature. 2006;443(7112):700–4.

17. Yemisci M, Gursoy-Ozdemir Y, Vural A, Can A, Topalkara K, Dalkara T. Pericyte contraction induced by oxidative-nitrative stress impairs capillary reflow despite successful opening of an occluded cerebral artery. Nat Med. 2009;15(9):1031–7.

18. Hall CN, Reynell C, Gesslein B, Hamilton NB, Mishra A, Sutherland BA, et al. Capillary pericytes regulate cerebral blood flow in health and disease. Nature. 2014;508(7494):55–60.

19. Alarcon-Martinez L, Yilmaz-Ozcan S, Yemisci M, Schallek J, Kilic K, Villafranca-Baughman D, et al. Retinal ischemia induces alpha-SMA-mediated capillary pericyte contraction coincident with perivascular glycogen depletion. Acta Neuropathol Commun. 2019;7(1):134.

20. Mishra A, Reynolds JP, Chen Y, Gourine AV, Rusakov DA, Attwell D. Astrocytes mediate neurovascular signaling to capillary pericytes but not to arterioles. Nat Neurosci. 2016;19(12):1619–27.

21. Kisler K, Nelson AR, Rege SV, Ramanathan A, Wang Y, Ahuja A, et al. Pericyte degeneration leads to neurovascular uncoupling and limits oxygen supply to brain. Nat Neurosci. 2017;20(3):406–16.

22. Swanson RA. Physiologic coupling of glial glycogen metabolism to neuronal activity in brain. Can J Physiol Pharmacol. 1992;70 Suppl:S138–44.

23. Pellerin L, Stolz M, Sorg O, Martin JL, Deschepper CF, Magistretti PJ. Regulation of energy metabolism by neurotransmitters in astrocytes in primary culture and in an immortalized cell line. Glia. 1997;21(1):74–83.

24. Zois CE, Harris AL. Glycogen metabolism has a key role in the cancer microenvironment and provides new targets for cancer therapy. J Mol Med (Berl). 2016;94(2):137–54.

25. Kilic K, Karatas H, Donmez-Demir B, Eren-Kocak E, Gursoy-Ozdemir Y, Can A, et al. Inadequate brain glycogen or sleep increases spreading depression susceptibility. Ann Neurol. 2018;83(1):61–73.

26. Duran J, Saez I, Gruart A, Guinovart JJ, Delgado-Garcia JM. Impairment in long-term memory formation and learning-dependent synaptic plasticity in mice lacking glycogen synthase in the brain. J Cereb Blood Flow Metab. 2013;33(4):550–6.

27. Hackett MJ, Sylvain NJ, Hou H, Caine S, Alaverdashvili M, Pushie MJ, et al. Concurrent Glycogen and Lactate Imaging with FTIR Spectroscopy To Spatially Localize Metabolic Parameters of the Glial Response Following Brain Ischemia. Anal Chem. 2016;88(22):10949–56.

28. Allaman I, Belanger M, Magistretti PJ. Astrocyte-neuron metabolic relationships: for better and for worse. Trends Neurosci. 2011;34(2):76–87.

29. Berthet C, Lei H, Thevenet J, Gruetter R, Magistretti PJ, Hirt L. Neuroprotective role of lactate after cerebral ischemia. J Cereb Blood Flow Metab. 2009;29(11):1780–9.

30. Kawasaki H, Kosugi I, Sakao-Suzuki M, Meguro S, Tsutsui Y, Iwashita T. Intracerebroventricular and Intravascular Injection of Viral Particles and Fluorescent Microbeads into the Neonatal Brain. J Vis Exp. 2016(113).

31. Karatas H, Erdener SE, Gursoy-Ozdemir Y, Gurer G, Soylemezoglu F, Dunn AK, et al. Thrombotic distal middle cerebral artery occlusion produced by topical FeCl(3) application: a novel model suitable for intravital microscopy and thrombolysis studies. J Cereb Blood Flow Metab. 2011;31(6):1452–60.

32. Dunn AK, Bolay H, Moskowitz MA, Boas DA. Dynamic imaging of cerebral blood flow using laser speckle. J Cereb Blood Flow Metab. 2001;21(3):195–201.

33. Swanson RA, Morton MT, Tsao-Wu G, Savalos RA, Davidson C, Sharp FR. A semiautomated method for measuring brain infarct volume. J Cereb Blood Flow Metab. 1990;10(2):290–3.

34. Walls AB, Sickmann HM, Brown A, Bouman SD, Ransom B, Schousboe A, et al. Characterization of 1,4-dideoxy-1,4-imino-d-arabinitol (DAB) as an inhibitor of brain glycogen shunt activity. J Neurochem. 2008;105(4):1462–70.

35. Bulmer D. Dimedone as an aldehyde blocking reagent to facilitate the histochemical demonstration of glycogen. Stain Technol. 1959;34(2):95–8.

36. Swanson RA, Sagar SM, Sharp FR. Regional brain glycogen stores and metabolism during complete global ischaemia. Neurol Res. 1989;11(1):24–8.

37. Wu L, Butler NJM, Swanson RA. Technical and Comparative Aspects of Brain Glycogen Metabolism. Adv Neurobiol. 2019;23:169–85.

38. Wu L, Wong CP, Swanson RA. Methodological considerations for studies of brain glycogen. J Neurosci Res. 2019;97(8):914–22.

39. Oe Y, Baba O, Ashida H, Nakamura KC, Hirase H. Glycogen distribution in the microwave-fixed mouse brain reveals heterogeneous astrocytic patterns. Glia. 2016;64(9):1532–45.

40. Subbarao KV, Stolzenburg JU, Hertz L. Pharmacological characteristics of potassium-induced, glycogenolysis in astrocytes. Neurosci Lett. 1995;196(1–2):45–8.

41. Pfeiffer-Guglielmi B, Francke M, Reichenbach A, Fleckenstein B, Jung G, Hamprecht B. Glycogen phosphorylase isozyme pattern in mammalian retinal Muller (glial) cells and in astrocytes of retina and optic nerve. Glia. 2005;49(1):84–95.

42. Dienel GA, Cruz NF. Astrocyte activation in working brain: energy supplied by minor substrates. Neurochem Int. 2006;48(6–7):586–95.

43. Cai Y, Guo H, Fan Z, Zhang X, Wu D, Tang W, et al. Glycogenolysis Is Crucial for Astrocytic Glycogen Accumulation and Brain Damage after Reperfusion in Ischemic Stroke. iScience. 2020;23(5):101136.

44. Perezleon JA, Osorio-Paz I, Francois L, Salceda R. Immunohistochemical localization of glycogen synthase and GSK3beta: control of glycogen content in retina. Neurochem Res. 2013;38(5):1063–9.

45. Suh SW, Bergher JP, Anderson CM, Treadway JL, Fosgerau K, Swanson RA. Astrocyte glycogen sustains neuronal activity during hypoglycemia: studies with the glycogen phosphorylase inhibitor CP-316,819 ([R-R*,S*]-5-chloro-N-[2-hydroxy-3-(methoxymethylamino)-3-oxo-1-(phenylmethyl)pro pyl]-1H-indole-2-carboxamide). J Pharmacol Exp Ther. 2007;321(1):45–50.

