## Supplementary Material for "Disrupted Cerebral Peri-Microvascular Glycogen Promotes Capillary Constrictions and Aggravates Ischemia in Mice"

**Supplementary Fig. 1**

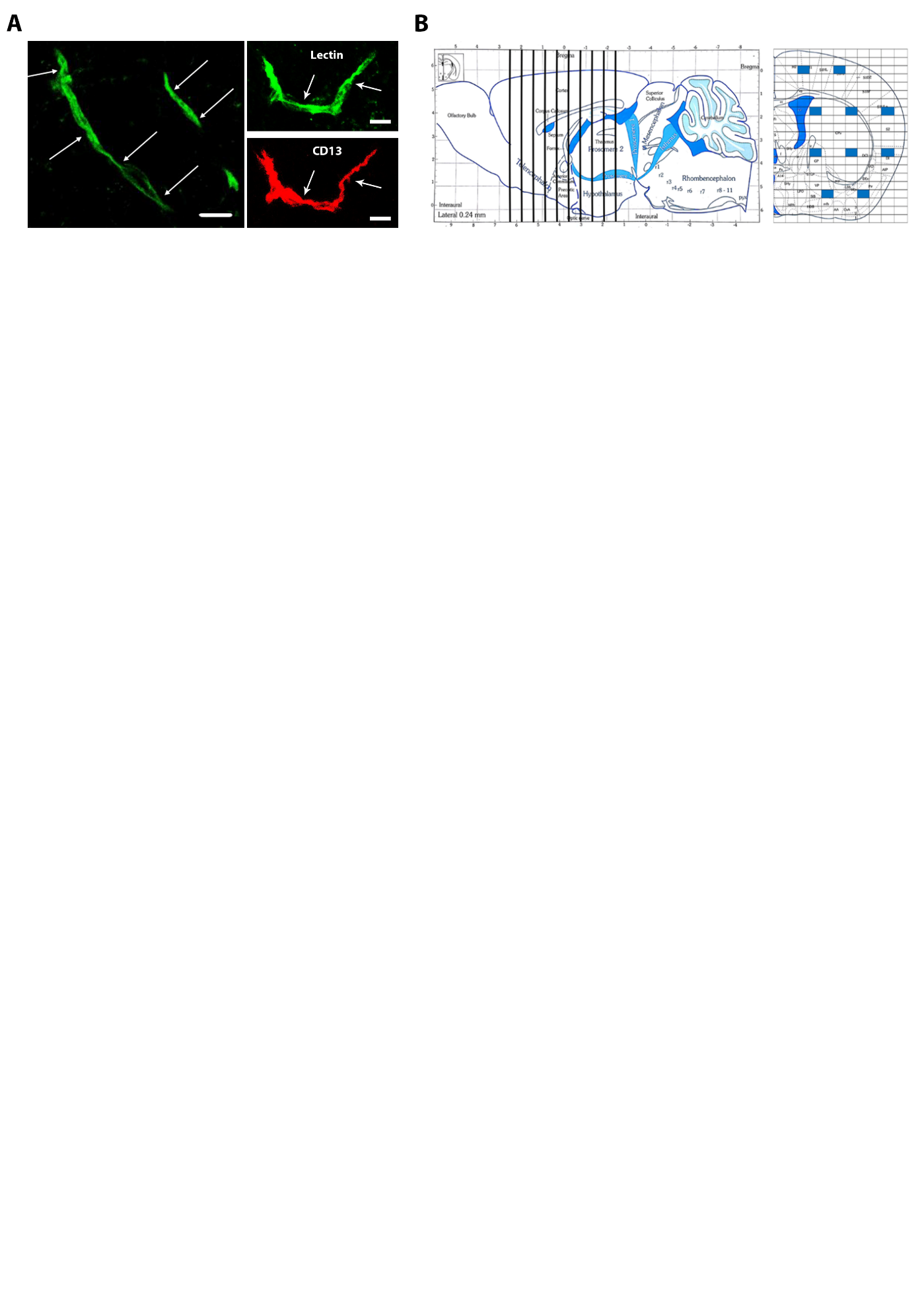

Quantification of Microvascular constrictions. (A) Double labelling with Lycopersicon Esculentum Lectin (left and right upper panels-green) and CD13 (right lower panel-red) demonstrates CD13+ pericyte mediated microvascular constrictions (arrows). Parameters defining constrictions are explained with further details in Methods. Images represent 3D reconstruction of 40-μm z-stack. Scale bars, 10 μm. (B) Illustrative images of mouse brains: sagittal view (left panel), coronal view (right panel). Modified from George Paxinos mouse brain atlas. Ten vertical black lines represent each coronal section taken for semi-stereological quantification (left panel). Vertical and horizontal black lines indicate each area at 40x magnification under fluorescent microscope (360 μm × 240 μm). Blue rectangles demonstrate the 10 areas in which quantification takes part (ROI size: 240 μm × 160 μm).

**Table 1.**

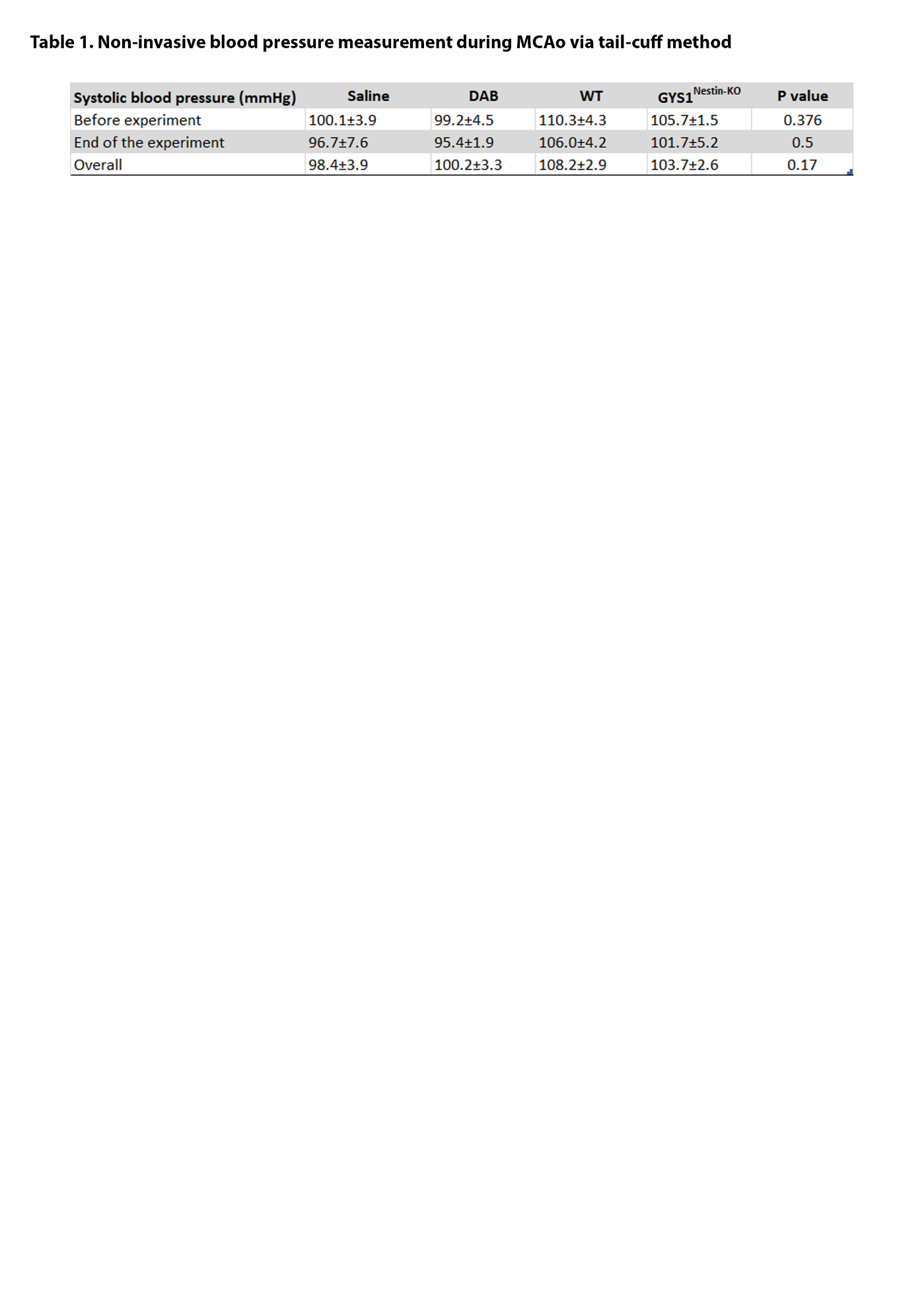

**Supplementary Fig. 2**

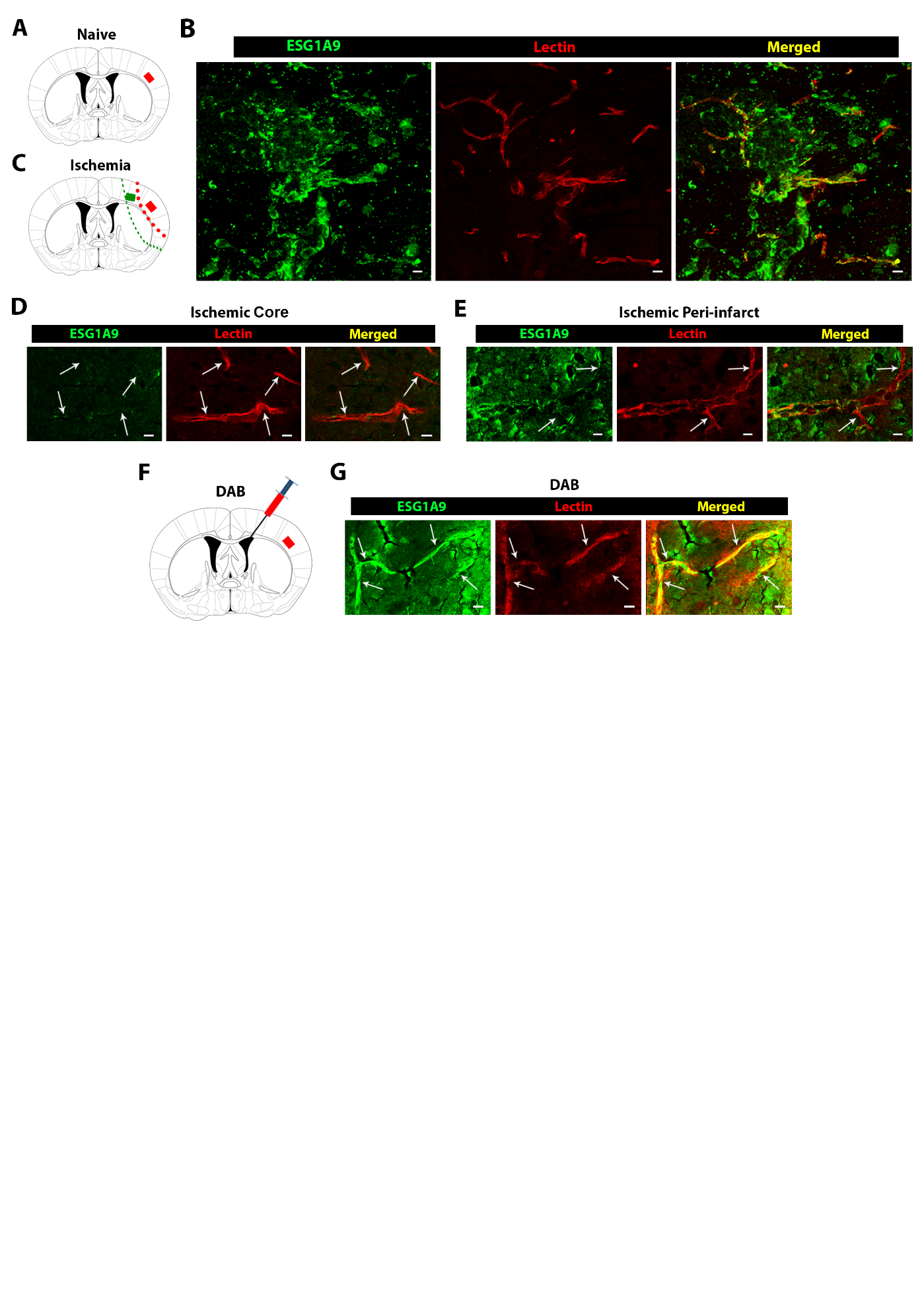

Immunofluorescence with anti-glycogen antibody ESG1A9 in naïve and ischemic mice. (A) Illustrative image of coronal section of mouse brain including MCA territory. Red box represents the area that images were taken. (B) Double labelling with ESG1A9 (left panel-green) and Lectin (middle panel-red) in naïve mice. Anti-glycogen antibody reveals substantial deposition around microvessels in merged image (right panel). Images represent 3D reconstruction of 40-μm z-stack. Scale bars, 10 μm. (C) Illustrative image of coronal section of mouse brain including MCA territory in ischemic mice. Red and green dotted lines represent the borders of core and peri-infarct, respectively. Also red and green boxes represent the core and peri-infarct ROIs in which images were taken, respectively. (D) Double labelling with ESG1A9 (green-left panel) and Lectin (red-middle panel) around core area in ischemic mice 2 hours after permanent MCAo. The loss of anti-glycogen antibody related signal intensity reveals the coincided nature of microvessel constrictions (arrows) and glycogen depletion (right panel). Images represent 3D reconstruction of 40-μm z-stack. Scale bars, 10 μm. (E) Double labelling with ESG1A9 (green-left panel) and Lectin (red-middle panel) around peri-infarct area in ischemic mice 2 hours after permanent MCAo. Anti-glycogen antibody related signal intensity increases in the parenchyma although the microvessel constrictions are observed where glycogen depletes (arrows). Images represent 3D reconstruction of 40-μm z-stack. Scale bars, 10 μm.
